# Time-Evolving Immunomodulatory Activity in Green-Fixed Devitalized Cellular Materials

**DOI:** 10.64898/2026.10.07.756712

**Authors:** Marlene C. Correia, Deniz Atila, Eduarda Mota-Silva, Mariana B. Oliveira

## Abstract

By decoupling biological function from cellular viability, fixed abiotic cellular materials have emerged as a promising platform for next-generation immunotherapies, preserving essential immunomodulatory activities while enabling unprecedented control over manufacturing, storage, and standardization. Nevertheless, the coupled effects of fabrication history and post-fabrication environmental cues on their biological functionality remain elusive. Here, we show that both preparation route and storage time critically affect the bioactivity of mesenchymal stromal cell (MSC)-derived microtissues. By exploring different devitalization agents, we demonstrate that an ethanol-based strategy provides a green fixation option with comparable preservation of volumetric and fixed cell architecture, as well as immunomodulatory function. Processing protocols did not merely substitute conventional fixation but tailored specific immune-related bioactivity. An unexpected processing protocol and time-dependent enhancement of immunomodulatory performance was observed, with ethanol-fixed microtissues maintaining a stable baseline bioactive profile while increasing functional potency over 8 weeks of storage. We establish a framework for the design of immunomodulatory abiotic cell-derived materials with potential for the treatment of immune-mediated diseases and regenerative therapies, and we hope that the principles uncovered here may contribute to a broader understanding of how devitalized tissues can be engineered, preserved, and functionally tuned, whether lab-made or derived from natural biological systems.

## Introduction

The preservation of human and animal tissues has long been sought for the biological understanding of animal and human biology, as well as for the preparation of one of the first diagnostic tools: histological sections [1,2]. By stabilizing tissue architecture and cellular features, fixation methods enabled the examination of tissues outside the living organism, with a major impact on anatomy, pathology, and experimental biology. From a therapeutic perspective, devitalized tissues have also been used in clinical practice, particularly in orthopedic reconstruction and bone grafting, including for the repair of defects generated by tumour resection [3]. The need to stabilize biological tissues is not exclusive to biology and medicine. In fact, even before significant advances were implemented in these fields, food preservation enabled remarkable progress in ways to preserve edible tissues [4]. Of historical note is the use of salt as a food preservative, capable of partially maintaining tissue structure, despite changes in texture, while preserving overall nutritional content and providing antimicrobial properties compatible with food safety [5,6].

Despite major advances in cell culture technologies and cell-based products, environmental sustainability has rarely been explored as a primary design consideration. Cellular microtissues assembled *in vitro* from human mesenchymal stromal cells (MSCs) and fixed with methanol preserved their macrostructure, as well as the original location of cell nuclei and several cell membrane proteins [7]. Their ability to interact with different immune cells, including monocytes, macrophages, and T lymphocytes, showed that, despite the lack of cellular viability, these constructs retain inherent biological activity [7]. Although methanol fixation has long been identified as an effective cell fixation method [8], its toxic nature requires protective measures for researchers preparing the samples, as well as specialized waste treatment with high environmental costs [9,10]. The use of less toxic solvents, such as ethanol, has also been reported for cell fixation [11,12]. However, its effectiveness for the preparation of devitalized tissues with preserved and comparable immunological properties remains unknown. Salts may represent a greener alternative for devitalized tissue preparation and storage. Inspired by traditional foods such as salted codfish and meat jerky, salts could be used as agents capable of promoting the drying of cellular microtissues, reducing water availability and supporting preservation without highly toxic organic solvents [5]. Of note, however, their effect on the preservation of biological domains, such as membrane-associated proteins and immune-interactive surfaces, is still poorly understood. The transition from methanol fixation to strategies using reagents and solvents with lower toxicity may enable the safer manufacturing of cell-derived devitalized products.

Not only the preparation but also the preservation of biological materials is critically important and remains a major practical and scientific challenge. Cellular, cell-derived (*e.g.*, proteins, extracellular vesicles, extracellular matrices) and other biologics (*e.g.*, recombinant RNAs) are often preserved through continuous exposure to cold settings, ranging from 4 °C to cryopreservation setting depending on the products [13]. Learning from century-old food preservation techniques could enable the long-term preservation of these products in cold chain-free settings, bringing cell therapies closer to environmentally sustainable practices. This is particularly relevant for advanced cell-derived materials, whose manufacturing, storage, and distribution are often limited by cost and logistics, often culminating in high environmental burden. Here, we explore alternative strategies for the preparation and preservation of devitalized cellular microtissues, namely ethanol fixation and salt-based drying, as potentially safer and greener approaches for advanced cell-derived materials with putative applications in regenerative medicine and immunomodulation. Beyond initial fixation outcomes, we also explore how distinct processing routes influence the temporal evolution of bioactivity during storage. We compare aqueous and dry preservation under refrigerated and ambient-compatible conditions over extended time periods, showing that preservation format and temperature exert only minor effects on structural and functional stability. This study establishes green processing and storage time as coupled, design-relevant parameters that define the functional landscape of devitalized cell-derived tissue constructs, offering a broader framework for the predictable engineering of abiotic cell-based systems, as well as for the design, preservation, and functional tuning of more complex materials derived from living matter.

## Results

To investigate whether the performance of devitalized cellular materials can be decoupled from hazardous processing and cold-chain requirements, we explored alternative strategies for their preparation and preservation. We developed alternative, lower-toxicity devitalization strategies compatible with ambient conditions, benchmarking them against our previously established methanol (MeOH)-based protocol [7].

Methodologies for cell aggregate fixation were based on the use of ethanol (EtOH) as a lower toxicity and less hazardous substitute for MeOH [14,15], and also on NaCl in different configurations as a simple, abundant, and biocompatible osmotic agent capable of promoting dehydration and molecular stabilization [5] (**Figure 1a**). Besides different fixation agents, different temperatures, either refrigerated (on ice) and room temperature (RT) were explored for the preparation of the devitalized materials. An initially tested condition was based on the use of 100% EtOH, administered to cell aggregates for 24h on ice, in a process that mirrored the optimized protocol using MeOH [7]. To further reduce reliance on refrigerated conditions and enable a more accessible and energy-efficient process, a stepwise optimization of RT-compatible conditions was subsequently undertaken. Among these, a formulation containing 70% EtOH (v/v) was explored based on the hypothesis that the presence of a controlled water fraction could moderate the rate of dehydration and facilitate more homogeneous solvent diffusion throughout the aggregates. This approach was expected to minimize the formation of a dense and stiff dehydrated shell at the aggregate periphery, a phenomenon commonly associated with rapid alcohol-induced dehydration at RT [16], which can hinder solvent penetration and compromise the preservation of internal architecture. NaCl-based conditions were further explored as reduced-solvent and solvent-free sustainable alternatives, including both near-saturated NaCl solution and solid salt-embedding approaches.

**Figure 1.**
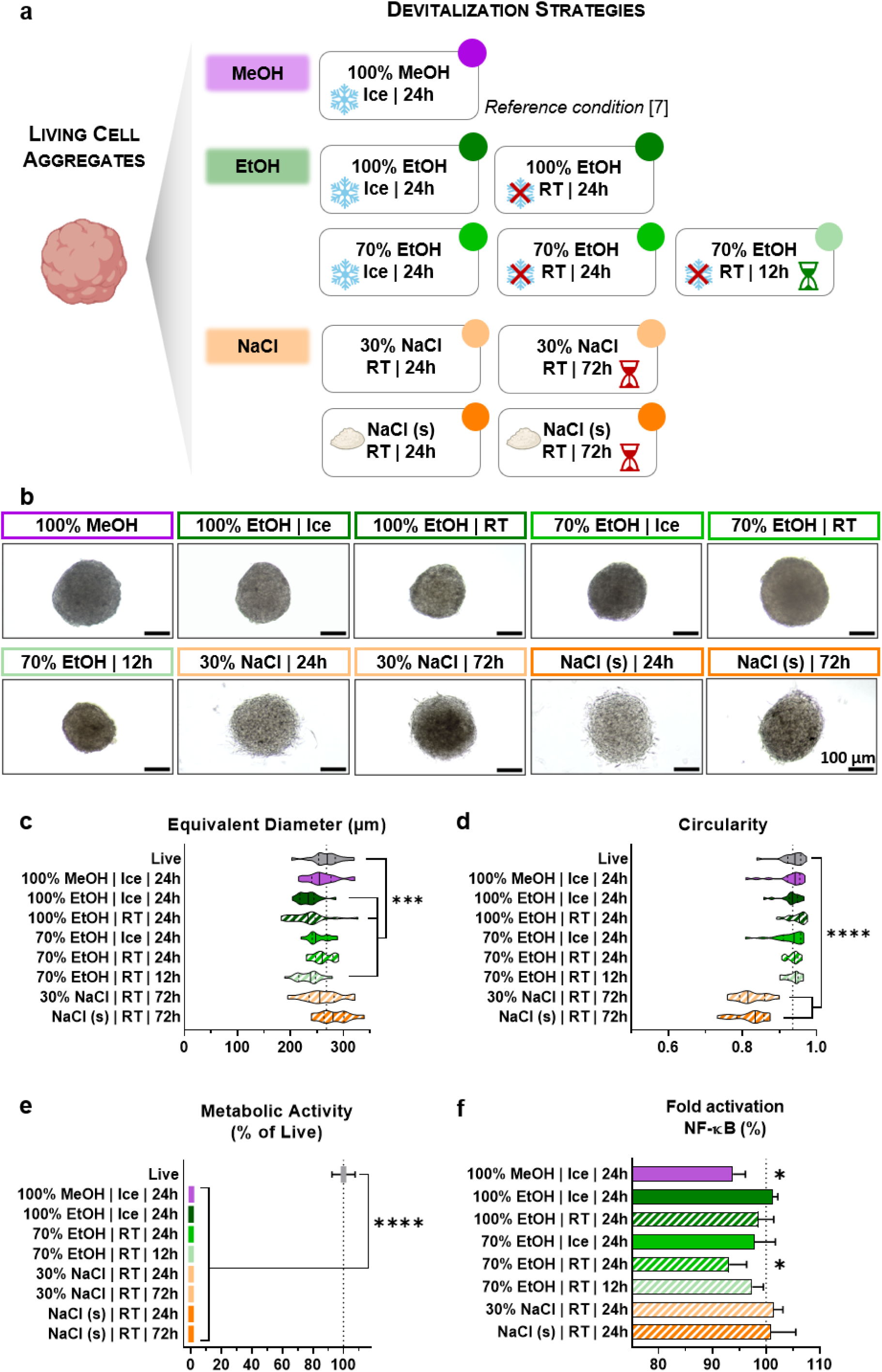
Screening and optimization of sustainable devitalization strategies. **a)** Schematic overview of the optimization workflow used to transition from the established methanol (MeOH)-based protocol toward simplified and more sustainable devitalization strategies. Ethanol (EtOH)-based approaches explored reduction of solvent toxicity and concentration, transition to RT processing, and shortened processing time; while NaCl-based conditions were investigated as reduced-solvent alternatives, including solvent-free NaCl embedding. **b)** Representative brightfield images of hASC spheroids following treatment under the indicated conditions. Scale bars: 100 µm. **c)** Equivalent diameter and **d)** circularity of live and devitalized spheroids quantified using the AnaSP software [20]. Data represent individual spheroids (n≈15 per condition) obtained from the same production batch. Statistical analysis was performed using one-way ANOVA with Dunnett’s multiple comparisons test against live spheroid control (*** p<0.001, **** p<0.0001). **e)** Relative metabolic activity of live and devitalized spheroids assessed by CellTiter-Glo®. All tested conditions resulted in a marked reduction in metabolic activity, confirming successful devitalization, including under NaCl-based conditions. Data are presented as mean ± SD. MeOH and all EtOH conditions were evaluated in two independent experiments using different donors (n=4-5 replicates per condition and experiment), whereas remaining screening conditions were assessed in one experiment (n=5 replicates per condition). Statistical analysis was performed using one-way ANOVA with Dunnett’s multiple comparisons test against the live control (****, p<0.0001). **f)** Early modulation of LPS-induced NF-κB activation in reporter monocytes (24h) by devitalized spheroids prepared under the indicated conditions, expressed relatively to the corresponding positive LPS-activated control within each experiment. Solid bars represent refrigerated processing whereas patterned bars show treatments performed at RT. Data are presented as mean ± SD from a single screening experiment (n=4 replicates per condition). Statistical analysis was performed by multiple unpaired t-tests against the corresponding positive control. Only the established MeOH protocol and the 70% EtOH RT 24h condition resulted in significant reduction in NF-κB activation (*, p<0.05).

Overall, alcohols-based devitalization produced only minor changes in spheroid morphology compared with live controls (**Figure 1b-d**). A slight tendency toward reduced equivalent diameter was observed, consistent with the dehydration associated with alcohol fixation, although statistically significant differences were limited to selected conditions. Circularity was largely preserved across alcohol-based protocols. Among the EtOH-based strategies, 100% EtOH induced a statistically significant reduction in equivalent diameter relative to live spheroids, particularly under RT processing, where increased variability was also observed. In contrast, the 70% EtOH treatment yielded morphometric characteristics comparable to the established MeOH condition under both ice and RT processing. Shortening the 70% EtOH treatment to 12h resulted in a significant reduction in equivalent diameter relative to live spheroids, while circularity remained largely unaffected. Initial NaCl-based treatments (for 24h) produced spheroids with poorly defined boundaries and reduced structural cohesion (**Figure 1b**), resulting in the release of cellular fragments during routine handling. Extending treatment to 72h slightly improved the mechanical integrity of the spheroids during handling. However, despite maintaining equivalent diameters close to those of live spheroids, circularity remained considerably lower than in the alcohol-devitalized conditions (mean circularity of 0.822 vs 0.928-0.947 for all other conditions). The devitalized character of all formulations was characterized through the measurement of metabolic activity after preparation. All tested conditions, including the shortened 70% EtOH 12h treatment, showed less than 0.2% metabolic activity compared to the live aggregate control (**Figure 1e**). Additionally, no increase in metabolic activity was observed after 7 days of culture in full medium (**Figure S1**), further confirming the absence of cellular regrowth and the irreversible nature of the devitalization process.

The different devitalization conditions were initially screened based on their capacity to modulate the NF-κB pathway of LPS-stimulated monocytes during the first 24h of exposure (**Figure 1f**). NF-κB is a central inflammatory signaling pathway activated by a wide range of stimuli, including cytokines, cellular stress signals, and microbial products, serving as a key regulator of innate immune responses [17]. Combined with morphometric quality analysis, this early functional assessment served to identify the most promising devitalization protocols, which were subsequently carried forward for more detailed bioactivity and preservation studies. As expected, control MeOH-devitalized spheroids significantly attenuated NF-κB activation, consistent with the retention of immunomodulatory activity following devitalization [7]. Among EtOH conditions, the EtOH 70% treatment for 24h at RT was the only other condition achieving significant reduction. NaCl-based conditions, besides showing structural instability, failed to reduce the activation level of NF-κB in reporter monocytes, suggesting their inadequacy as predictable and functional immunomodulatory devices. The combined evaluation of structural integrity, functional activity, and the advantages of RT processing led to the selection of 70% EtOH, 24h RT condition (hereafter referred to as EtOH) for subsequent studies. Additional characterization by Live/Dead staining confirmed effective devitalization under the selected EtOH protocol relative to the live spheroid control, and histological characterization confirmed structural preservation compared to traditional paraformaldehyde-fixed and control MeOH-treated spheroids (**Figure S2**).

Immunomodulatory activity following devitalization was further evaluated using NF-κB and cGAS-STING reporter monocyte assays for longer actuation periods, up to 72h (**Figure 2a**). cGAS-STING was additionally assessed here as a complementary innate immune pathway that detects cytosolic double-stranded DNA and triggers downstream interferon-associated and inflammatory responses [18]. To probe the reproducibility of the immunomodulatory response, assays were performed using devitalized aggregates generated from three independent production batches and from hASCs derived from two independent donors. EtOH-devitalized aggregates exhibited responses comparable to those observed with MeOH-treated materials. At 72 h, a sustained and more pronounced reduction in both NF-κB and cGAS–STING activation was observed in monocytes stimulated with LPS or cGAMP, respectively. In contrast, effects at 24 h were modest and did not reach statistical significance, likely reflecting inter-donor variability and early-stage response dynamics. The attenuation of the cGAS-STING axis is particularly relevant here, as fragmented DNA remnants generated during devitalization could otherwise act as damage-associated signals, capable of stimulating innate immune activation [18]. Its suppression suggests that devitalized hASC-aggregates, prepared with either fixative, do not elicit a significant DNA-driven innate immune response despite the presence of cellular remnants. These findings are consistent with previous work [7], indicating that MeOH-based devitalization preserves key bioactive immunomodulatory cues. Collectively, this supports that the optimized EtOH fixation protocol similarly maintains functional immunomodulatory components within the extracellular matrix and cellular remnants, enabling sustained pathway attenuation across independent donor-derived hASC preparations. To further evaluate the versatility of the optimized EtOH protocol, devitalization was applied to millimeter-scale hASC-derived tissue constructs with distinct architectures (**Figure 2b**). Patterned membrane constructs preserved their macroscopic organization following devitalization, while ring-shaped constructs retained sufficient structural integrity to undergo manual stretching after devitalization. These results extend the applicability of the protocol beyond spheroid formulations, supporting its use across engineered tissue constructs of varying sizes and complexity.

**Figure 2.**
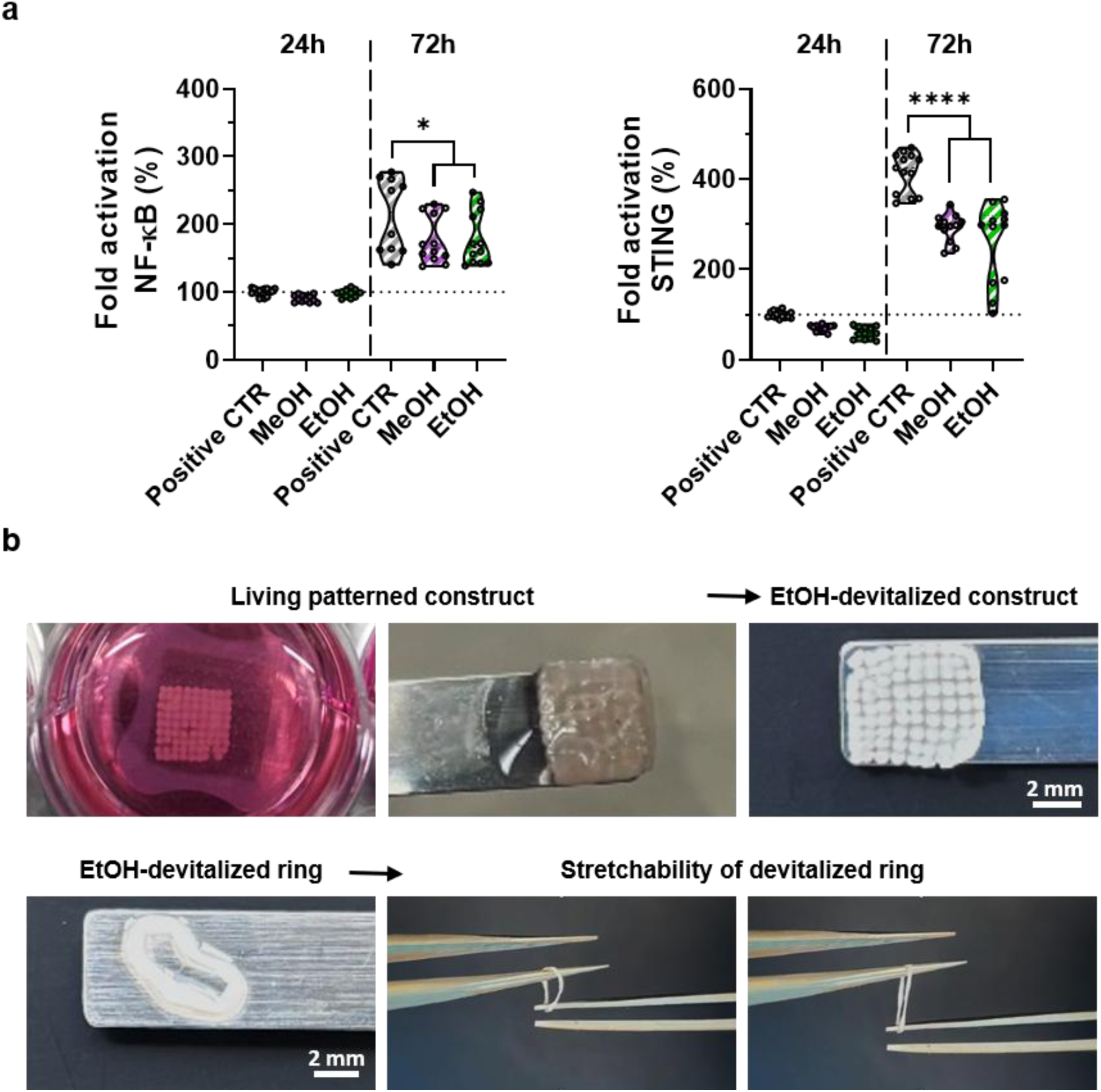
Structural and functional validation of the optimized ethanol devitalization protocol. **a)** Modulation of NF-κB and cGAS-STING activation following exposure to MeOH- and EtOH-devitalized spheroids, evaluated at early (24h; solid violins) and sustained (72h; patterned violins) timepoints. The 70% EtOH RT 24h condition (referred as EtOH) selected from the screening workflow was compared directly with the established MeOH protocol. Responses were normalized to the corresponding positive control at 24h. Violin plots represent data distribution across three independent experiments using independent spheroid batches produced from two different donors (total of 10-14 replicates per condition across experiments). Statistical analysis was performed using one-way ANOVA with Dunnet’s multiple comparisons test against the corresponding positive control (** p<0.01, *** p<0.001, **** p<0.0001). **b)** Representative images of millimeter-scale hASC-derived tissue constructs following devitalization using the optimized 70% EtOH RT protocol. Patterned membrane construct retained their overall geometry following devitalization, while ring-shaped constructs remained sufficiently cohesive for manual stretching. Scale bars: 2 mm.

Preservation represents a key requirement for the practical translation of devitalized cellular materials, as their utility depends not only on the retention of structural integrity but also on the maintenance of biological activity beyond the time of fabrication. In previous work, MeOH-fixed spheroids preserved their ability to attenuate NF-κB activation following storage in DPBS at 4 °C for up to 2 months [7]. Building on these observations, we investigated whether more sustainable preservation strategies could support long-term functionality while reducing reliance on aqueous media and refrigerated conditions (**Figure 3a**). The first step relied on exclusion of refrigerated requirements, with MeOH- or EtOH-fixed samples stored in sterile DPBS for a course of 8 weeks at RT. Similar morphometric trends were observed for MeOH- and EtOH-devitalized spheroids stored in DPBS at both temperatures (**Figure 3b-c**). An initial increase in equivalent diameter relative to pre-storage values was observed after 1 week, followed by a tendency toward progressively reduced diameter. This increase may reflect rehydration following alcohol devitalization. Circularity also remained slightly above pre-storage values across all aqueous storage conditions and timepoints, indicating largely preserved spheroid shape throughout storage at both temperatures. Dry preservation methodologies were also explored, as they may enable ambient-compatible storage while offering practical advantages for the deployment of devitalized cellular materials. Removal of the liquid phase would simplify handling and transportation, reduce contamination risks, while being compatible with a decrease in dependence on refrigerated infrastructure. From a manufacturing perspective, dry materials may also facilitate integration into more complex multi-material systems, such as scaffolds, coatings, and other composite systems. Among the drying-based approaches, air-drying (at RT, in a laminar flow chamber, overnight) was compared with freeze-drying after MeOH-devitalization (**Figure S3**). When compared to storage in DPBS, both drying methods led to a reduction of the equivalent diameter of devitalized aggregates following rehydration in DPBS for 1 day. Freeze-dried samples exhibited altered aspect ratio, reduced compactness, and visibly distorted morphologies upon rehydration, indicating a more pronounced disruption of spheroid morphology. As air-drying provided improved structural preservation while relying on a simpler preservation protocol, freeze-drying was not investigated further, and subsequent studies focused on air-dried materials. In addition to aqueous and air-dried preservation, we investigated storage in solid NaCl as a solvent-free strategy inspired by conventional dehydration approaches and the ability of highly concentrated salt environments to reduce water activity. Dry preservation largely maintained spheroid diameter following rehydration (**Figure 3b**). Significant deviations from the pre-storage condition were observed only for MeOH-devitalized air-dried spheroids at 3 weeks. Equivalent diameter of EtOH-devitalized spheroids did not differ significantly from pre-storage values throughout the preservation period. In contrast, circularity exhibited a clearer time-dependent response (**Figure 3c**). Individual air-dried spheroids maintained their circularity after rehydration following short-term storage (1 week), whereas significant reductions were observed after 3 and 8 weeks for EtOH-devitalized spheroids, indicating progressively less circular spheroid morphology following air-drying and prolonged storage. A fraction of air-dried spheroids formed cohesive spheroid assemblies during preservation (**Figure S4**). Following rehydration, pipetting enabled recovery of individual spheroids, although a part of the aggregates remained as small, interconnected assemblies, limiting the number of spheroids that could be accurately segmented for automated morphometric analysis.

**Figure 3.**
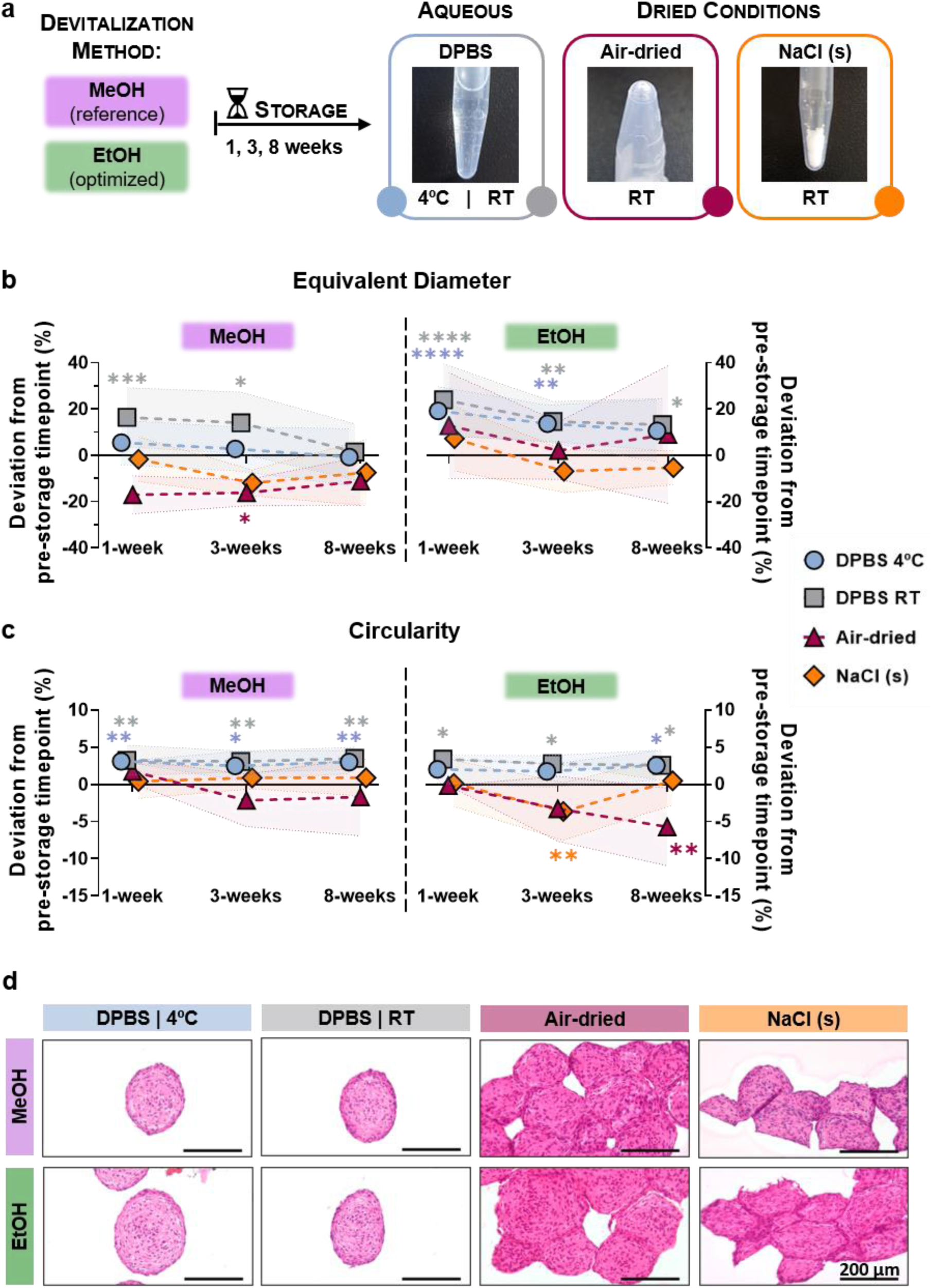
Structural stability of devitalized spheroids during long-term preservation. **a)** Schematic overview of the preservation strategies evaluated following selection of the optimized EtOH devitalization condition, in comparison with the established MeOH protocol. Aqueous preservation included DPBS storage at 4 °C and RT, while dry preservation strategies comprised air-drying and NaCl embedding. Samples were stored for up to 8 weeks prior to characterization. **b)** Equivalent diameter and **c)** circularity of MeOH-(left) and EtOH-devitalized (right) spheroids during preservation, expressed as percentage deviation from the corresponding pre-storage condition (Figure 1c-d) after 1, 3, and 8 weeks of storage. For dry preservation conditions, measurements were performed following rehydration in DPBS and handling to separate dried assemblies. Symbols represent mean values at each timepoint for each preservation condition, connected across the preservation period, through a dashed line, with shaded regions indicating SD. Data were obtained from a single spheroid production batch. Due to increased aggregation and contact between spheroids following dry preservation, fewer individual spheroids could be reliably segmented for AnaSP analysis in air-dried and NaCl-embedded conditions compared to aqueous preservation conditions (n=12-25 for aqueous conditions and n=2-12 for dry conditions). Statistical analysis was performed using one-way ANOVA with Dunnett’s multiple comparisons test against the corresponding pre-storage condition (* p<0.05, ** p<0.01). **d)** Representative H&E-stained histological sections of MeOH- and EtOH-devitalized spheroids following 8 weeks of storage under aqueous and dry preservation conditions. For dry preservation conditions (air-dried and NaCl-embedded), no attempt to separate spheroid assemblies was performed prior to analysis in order to preserve the structures formed during storage. Scale bars: 200 µm.

Histological analysis after 8 weeks of storage showed preservation of the overall spheroid architecture across all conditions (**Figure 3d**). Under aqueous storage (DPBS at 4 °C and RT), both MeOH- and EtOH-devitalized materials maintained their overall morphology with well-defined nuclei observed throughout the spheroid sections. Aggregates preserved using dry preservation strategies were analysed following rapid rehydration and without prior pipetting-based dissociation, therefore appearing as assembled spheroid structures reflecting the organization established during storage. As individual spheroids within the assembly are sectioned at different depths, their appearance in histological section may vary depending on the sectioning plane. Nevertheless, H&E staining showed comparable cellular organization and nuclear staining to that observed under aqueous conditions. Individual spheroids remained identifiable within the dry-preserved assemblies, although MeOH-devitalized samples showed more clearly defined and regular spheroid boundaries than the corresponding EtOH-devitalized materials following histological processing.

The biological functional performance of preserved devitalized spheroids was evaluated using NF-κB and cGAS-STING reporter monocyte assays, with responses expressed relative to the corresponding pre-storage timepoint (**Figure 4a-b**). For the NF-κB pathway, MeOH-devitalized spheroids largely maintained their immunomodulatory activity throughout the 8-week preservation period across all storage conditions, with spheroids embedded in NaCl for 8 weeks showing even an enhanced attenuation compared to the pre-storage response. EtOH-devitalized spheroids similarly preserved their immunomodulatory activity, while showing also un unexpected significant enhanced suppression of NF-κB activation following storage for 8 weeks. For the immediate response (24h), this enhancement was significantly pronounced after 8 weeks, when DPBS at RT and NaCl storage showed an average 23-42% greater attenuation than the corresponding pre-storage condition. The sustained response (72h) showed a transient enhancement after 3 weeks across dry preservation conditions for MeOH-devitalized spheroids and for all preservation strategies for EtOH-devitalized materials, returning to values comparable to the pre-storage response after 8 weeks. Preservation affected cGAS-STING modulation to a greater extent than NF-κB, with reductions in suppressive activity observed relative to the pre-storage condition (**Figure 4b**). For the immediate response (24h), both MeOH- and EtOH-devitalized spheroids exhibited significant reduced attenuation following storage across all conditions, with an apparent recovery towards the pre-storage response after 8 weeks. At the sustained timepoint (72h), MeOH-devitalized spheroids largely maintained their suppressive activity throughout storage, whereas EtOH-devitalized spheroids showed a modest but significant reduction in attenuation from their corresponding pre-storage response, although this difference became less pronounced after 8 weeks of storage. Factorial analysis further indicated that storage time accounted for a larger contribution of the observed functional variability than either devitalization strategy or preservation conditions (**Figure S5**), suggesting that those factors exerted a minor influence on pathway modulation.

**Figure 4.**
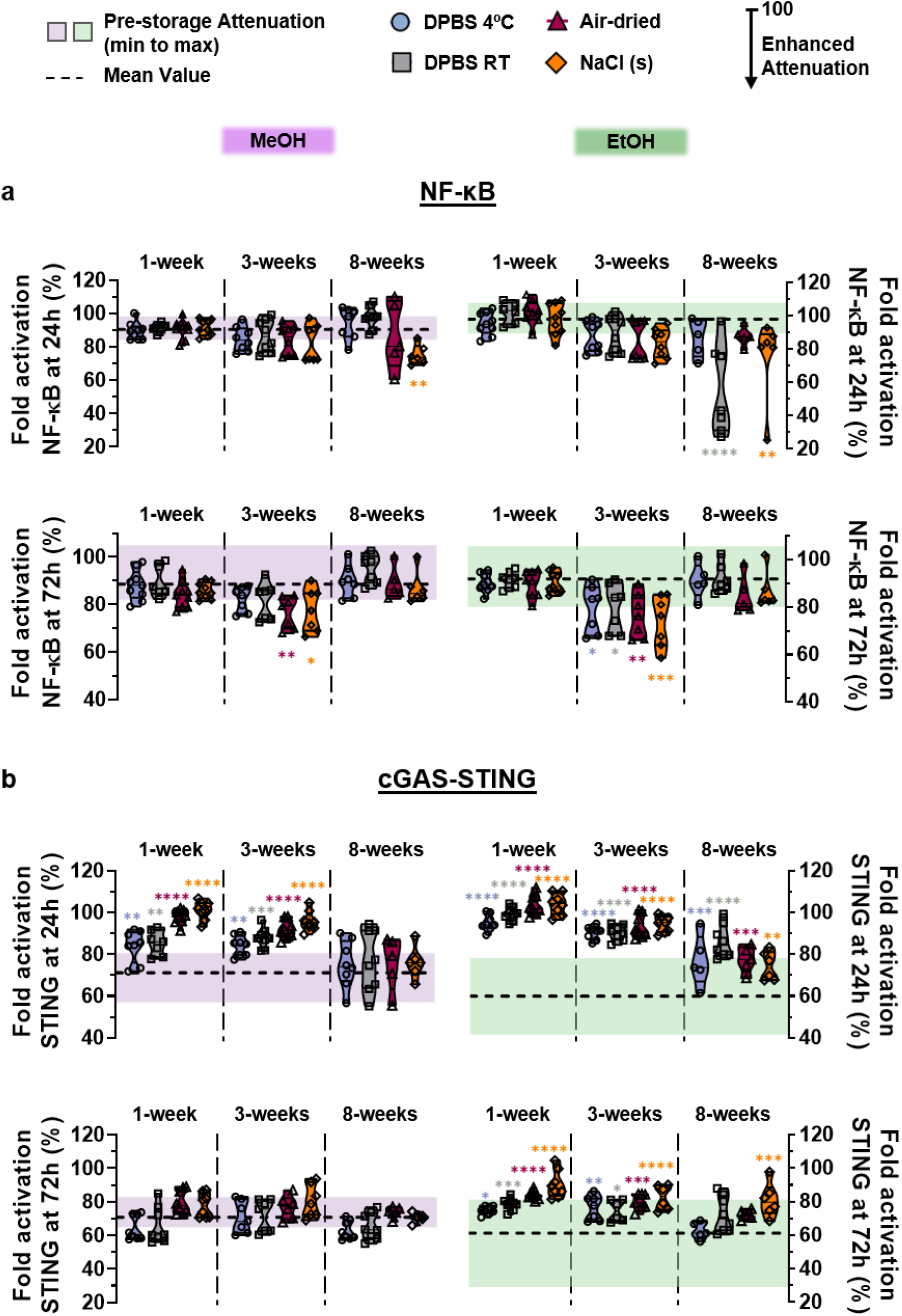
Functional stability of devitalized spheroids during long-term preservation. **a)** NF-κB and **b)** cGAS-STING modulation following exposure to preserved devitalized spheroids. Early (24h) and sustained (72h) responses are shown for MeOH-devitalized spheroids (left) and EtOH-devitalized spheroids (right). Responses are expressed relative to the corresponding positive control within each experiment; and the corresponding pre-storage attenuation results (Figure 2a) are shown by the shaded region in each graph. Monocyte activation was induced using LPS (for NF-κB) or cGAMP (for cGAS-STING). Devitalized spheroids were preserved under aqueous conditions (DPBS at 4°C or RT) or dry preservation approaches (air-drying and NaCl embedding) for 1, 3, and 8 weeks. Statistical analysis was performed using one-way ANOVA with Dunnett’s multiple comparisons test relative to the pre-storage timepoint (* p<0.05, ** p<0.01, *** p<0.001, **** p<0.0001). Violin plots represent data distribution across 2-3 independent experiments using independent spheroid batches produced from different cell donors (total of 5-9 replicates per condition across experiments).

## Discussion

Cellular technologies have enabled significant advances in biotechnology, regenerative medicine, and therapeutic development. While considerable efforts have focused on improving their biological performance and translational potential, less attention has been given to the sustainability of the processes required to produce, preserve, and distribute them. Identifying strategies that maintain functionality while reducing resource intensity may help bridge performance and sustainability considerations in next-generation biomaterial platforms. Beyond achieving biological function, understanding how process design, preservation requirements, and environmental impact collectively influence the development of cellular and cell-derived products may enable the predictive platform of a new generation of cell- and tissue-based materials.

Devitalized cellular materials provide an opportunity to align aspects of cellular technologies with resource-conscious and greener processing strategies. By converting living cellular constructs into stabilized abiotic bioactive materials, functional characteristics can be retained while decoupling performance from the constraints associated with maintaining viable cell systems [3]. While being effective at preserving structural integrity, MeOH introduces relevant safety and handling constraints due to its well-known toxicity [9,10]. Its use typically requires controlled laboratory conditions, specific waste management procedures, and restricted disposal pathways. Despite its chemical similarity to MeOH, EtOH presents a more favorable safety profile. In addition, EtOH can be derived from renewable fermentation processes [19], which may offer advantages in terms of resource origin depending on production routes. Importantly, the main difference between EtOH and MeOH in this context is not necessarily reflected in large differences in direct carbon footprint at laboratory scale, but rather in downstream implications related to hazard classification, handling requirements, and waste treatment complexity. Experimentally, both near-saturated solutions and solid-state salt conditions were ineffective in preserving the structural integrity of the devitalized materials under the conditions tested, indicating that salt-mediated stabilization alone is insufficient to maintain the preserved cellular architecture.

Storage conditions represent another relevant factor influencing overall resource demand for cellular material workflows. A simplified comparison between refrigerated storage at 4 °C and RT storage indicates a consistent difference in energy demand associated with continuous cooling requirements. Refrigerated storage relies on uninterrupted energy input to maintain temperature stability, whereas RT storage operates with minimal active energy consumption beyond baseline laboratory conditions. Although aqueous RT storage demonstrated that refrigeration may not be required to maintain the structural and functional properties of devitalized spheroids, dry preservation provides additional logistical advantages. By eliminating the liquid phase, dry-preserved materials may be easier to store, transport, and distribute, while reducing contamination risks and simplifying handling requirements. Dry preservation resulted in the formation of assembled spheroid structures that persisted after rehydration, although recovery of individual spheroids was possible through mechanical handling. While this feature limited automated morphometric analysis, it does not necessarily represent a limitation for downstream applications, where complete dissociation into individual spheroids may not be required. Rather, the assembled state generated during drying may represent an alternative material format.

Of particular interest, the biological performance of devitalized microtissues was not static throughout preservation. While preservation studies are typically designed to determine whether functionality can be maintained over time, our results suggest that storage duration itself can influence biological output. NF-κB attenuation remained preserved throughout storage and, in EtOH-devitalized materials, became more pronounced after 8 weeks. Similarly, cGAS-STING responses, which exhibited reduced attenuation at earlier timepoints, showed a tendency to recover towards pre-storage levels at later stages. These observations indicate that distinct processing routes generate materials that follow different functional trajectories during preservation, with EtOH fixation supporting a progressive enhancement of immunomodulatory activity over time. Although the mechanisms underlying these changes remain unclear, they suggest that preservation is not merely a passive process that maintains a predefined biological state. Instead, storage may influence the presence, accessibility, or release of bioactive cellular components retained within the devitalized material. In this context, time emerges as an active parameter, capable of shaping biological performance, rather than simply a measure of stability. This perspective expands the role of preservation from maintaining function to potentially tuning function, introducing temporal evolution as a previously underappreciated design dimension for devitalized cellular materials. Future studies investigating the molecular basis of these time-dependent effects may provide new opportunities to engineer materials whose biological activity can be programmed through both processing and storage history.

From a sustainability perspective, the combination of EtOH-based devitalization and ambient-compatible preservation strategies reduces reliance on hazardous solvents, refrigerated processing, and cold-chain storage. Beyond these practical advantages, our findings suggest that processing methodology and storage history jointly define the functional behavior of devitalized cellular materials. By demonstrating that biological activity can evolve over time as a function of preparation route, this work introduces a broader framework in which performance is not solely determined at the point of fabrication but continues to develop throughout preservation. We anticipate that this concept may contribute to the design of next-generation cell-derived materials and devitalized biological systems, where processing, preservation, and temporal evolution are considered together as interconnected variables governing long-term functionality. Such a perspective may be relevant not only for immunomodulatory biomaterials and regenerative medicine, but also for the broader engineering of living-derived constructs, biological interfaces, and stabilized bioactive systems.

## Conclusion

We demonstrate that the production and preservation of devitalized cellular materials can be substantially simplified without compromising structural or functional performance. We show that EtOH-based devitalization effectively inactivates and preserves structural and functional properties of hASC spheroids comparable to the previously established MeOH protocol, while enabling RT processing and reducing reliance on toxic solvents. In addition, we show that both aqueous and dry preservation strategies support the long-term stability of both MeOH- and EtOH-fixed constructs for up to 8 weeks. While aqueous storage remains effective, dry approaches such as air-drying and salt-embedding offer clear advantages by enabling storage in a non-aqueous state and reducing energy demands associated with refrigeration. Importantly, devitalized spheroids maintained their structural integrity and immunomodulatory function over time, and in some cases, enhanced activity after storage periods. Together, these findings establish a practical framework for the development of shelf-stable, cell-derived materials produced through lower-energy demanding and more accessible processing protocols. By aligning fabrication and preservation strategies with principles of sustainability and scalability, this work supports the broader translation of devitalized cellular materials as accessible and robust platforms for advanced bioactive applications.

## Experimental Section

### Cell Culture

Human adipose-derived mesenchymal stem/stromal cells (hASCs) were obtained from Lonza (PT-5006, lot no. 22TL108499 and 22TL018258), RoosterBio (C46001AD, lot no. 210323) and PromoCell (C-12977, lot no. 485Z035). Cells were expanded in T-flasks using α-minimum essential media (α-MEM; Gibco) supplemented with 0.22% (w/v) sodium bicarbonate (Sigma-Aldrich), 10% (v/v) heat-inactivated fetal bovine serum (FBS; Gibco) and 1% (v/v) antibiotic-antimycotic (100X stock, Gibco). Upon reaching confluence, cells were rinsed with Dulbecco’s Phosphate-Buffered Saline (DPBS, Sigma-Aldrich) and detached using 0.25% (v/v) trypsin-EDTA (Gibco). Devitalized materials were prepared from hASCs at passages 6-8. Cultures were maintained at 37°C in a humidified 5% CO_2_ incubator, with medium changes every 2-3 days.

THP1-Lucia™ NF-κB reporter cells (InvivoGen, thpl-nfkbv2, lot no. T13bis-4601) and THP1-Dual™ NF-κB/cGAS-STING reporter cells (InvivoGen, thpd-nfis, lot no. T14-5501) were used for immunomodulatory assays. Cells were cultured in Roswell Park Memorial Institute (RPMI) 1640 medium (Gibco) supplemented with 10% (v/v) heat-inactivated FBS, 1% (v/v) penicillin-streptomycin (100X stock, Gibco), 25 mM HEPES buffer (Sigma-Aldrich) and 100 µg mL^-1^ Normocin® (InvivoGen). After thawing, cells were initially cultured in medium containing 20% FBS to allow recovery, followed by gradual adaptation to standard 10% FBS conditions. After approximately two passages, antibiotic selection was reintroduced to maintain reporter stability. THP1-Lucia™ NF-κB culture was supplemented with 100 µg mL^-1^ Zeocin®, whereas THP1-Dual™ cells were cultured with 100 µg mL^-1^ Zeocin® and 10 µg mL^-1^ Blasticidin. Cells were maintained at 37°C in a humidified 5% CO_2_ incubator at a density of 0.4-0.5 x 10^6^ cells mL^-1^.

### Fabrication and Storage of Devitalized Materials

Spheroids were prepared using 2% (w/v) agarose micro-wells produced with 3D Petri Dish® molds (Sigma-Aldrich, Z764019), following manufacturer’s instructions. hASCs were seeded at a density corresponding to 4,000 cells per spheroid and allowed to aggregate for 48h. Following aggregation, spheroids were devitalized by incubation in MeOH [7], 100% EtOH or 70% EtOH. MeOH and 100% EtOH treatments were performed on ice, whereas 70% EtOH treatment was carried out at RT. Incubation was conducted for 24h under gentle agitation (20 rpm) on a rocking shaker, and a shortened 12h incubation in 70% EtOH was evaluated as an attempt to reduce processing time. Moreover, various geometric constructs could be prepared out of devitalized hACSs using agarose molds for aggregation. Patterned membranes and continuous rings were formed upon seeding 4 × 10^6 and 1 × 10^6 cells per construct, respectively. Once the intended geometries were observed, subsequent structures were removed from cell culture media, immersed in 70% EtOH, and devitalized for 24h at RT under gentle agitation.

For air-drying preservation, most of the devitalizing solution was removed and spheroids were allowed to dry under sterile conditions in a laminar flow cabinet. Dried spheroids were subsequently stored at RT for 1, 3, or 8 weeks. For other preservation conditions, devitalized spheroids were washed in DPBS for 24h at RT under the same agitation condition to remove residual devitalizing agents. For aqueous preservation, spheroids were maintained in DPBS at either 4°C or RT for the same storage periods. For NaCl embedding, washed spheroids were embedded in solid NaCl and stored at RT in a desiccator. Prior to use, NaCl was dissolved in sterile water to recover spheroids, followed by washing in 70% EtOH to ensure sterility.

Additional exploratory conditions, including NaCl-based devitalization (using 30% NaCl solution or solid NaCl) and freeze-drying preservation (-90°C, 0.0005 mbar, 24h) were initially evaluated but were not pursued further due to suboptimal structural integrity of devitalized spheroids. In the exploratory freeze-drying experiment, spheroids were produced using 8,000 cells per spheroid.

### Morphometric characterization

Brightfield images of spheroids were acquired before and after devitalization, as well as after 1, 3, and 8 weeks of preservation, using a Zeiss Primovert microscope equipped with an Axiocam 208c camera and ZEN software. For each condition, multiple spheroids were analysed from distinct production batches and/or cell donors, except for exploratory conditions reported in the SI. Images were converted to 8-bit grayscale in ImageJ for standardized processing. Morphometric characterization was performed using the automated AnaSP software [21]. Parameters extracted included equivalent diameter, circularity, Feret aspect ratio, and compactness, providing quantitative measures of spheroid size, roundness, elongation, and structural integrity, respectively. Based on the intended analysis, specific parameters were selected for comparative analysis.

### Metabolic activity measurement

The efficacy of the devitalization procedures was assessed using the 3D Cell Viability CellTiter-Glo® assay (Promega). Devitalization spheroids were compared to living spheroids used as a positive control. Measurements were performed immediately after devitalization, and after 7 days of culture in complete media under standard incubator conditions to evaluate potential regrowth. Two independent experiments were conducted using different cell donors, with five replicates per condition in each experiment.

For each condition, 10 spheroids suspended in α-MEM were incubated with CellTiter-Glo® reagent at a 1:1 ratio for 60 min at RT under agitation on a rocking shaker (80 rpm), protected from light. A mixture of CellTiter-Glo® reagent and α-MEM (1:1) was used as the blank control. Following incubation, plate was left undisturbed for 10 minutes to stabilize the signal, according to the manufacturer’s instructions. Luminescence was measured using a Varioskan Lux microplate reader (Thermo Fisher Scientific) in white opaque 96-well plates.

### Histologic analysis

For morphological analysis, spheroids were fixed in 4% formaldehyde (Sigma-Aldrich), dehydrated through a graded EtOH series, cleared, embedded in paraffin, and sectioned at 6 µm thickness. Sections were stained with a Hematoxylin and Eosin (H&E) kit (Bio-Optica, Italy) for general histological evaluation, following the manufacturer’s instructions. H&E-stained sections were mounted and imaged under brightfield microscopy.

### NF-κB and cGAS-STING pathway suppression following contact with devitalized spheroids

Assays were performed using complete RPMI 1640 medium, but without Normocin®, Zeocin® or Blasticidin. For each condition, 12 devitalized spheroids per replicate were added to THP-1 reporter cells, and all experimental conditions were performed in quadruplicate. Cultures were maintained at 37°C in a humidified incubator with 5% CO_2_. Positive controls (defined as 100% activation) consisted of reporter cells stimulated in the absence of spheroids: 100 ng mL^-1^ lipopolysaccharide (LPS, InvivoGen) for THP1-Lucia^TM^ NF-κB cells and 20 µg mL^-1^ 2’3’-cGAMP (InvivoGen) for THP1-Dual^TM^ cells, each performed in five replicates. A blank control (cells cultured in RPMI only, without spheroids or stimuli) was also included in five replicates in every experiment.

To evaluate the immunomodulatory capacity of the devitalized spheroids, reporter cells were cultured with spheroids in the presence of the corresponding positive stimulus (LPS or cGAMP) and compared to the respective positive control. To confirm that spheroids alone did not trigger pathway activation, at least one replicate of basal activation control was included for each spheroid condition (12 spheroids, no stimulus). Luminescence was measured after 24 h and 72 h of incubation, using culture supernatant mixed with QUANTI-Luc^TM^ 4 Reagent (InvivoGen) in white-opaque flat-bottom 96-well plates in a Varioskan Lux microplate reader (Thermo Fisher Scientific). In an exploratory screening, NF-κB modulation was also assessed using the THP1-Dual^TM^ cell line by measuring the absorbance at 620 nm in a Synergy HTX microplate reader (Biotek) following incubation of culture supernatant with QUANTI-Blue^TM^ Solution (InvivoGen).

### Statistical and factorial analysis

Statistical analyses were performed using GraphPad Prism version 8.0.2. Data were normalized as described in the corresponding figure captions. Differences between groups were assessed using multiple unpaired t-tests or one-way ANOVA followed by Dunnett’s multiple comparisons tests, as specified in the respective figure captions. Statistical significance was defined as * p<0.05, ** p<0.01, *** p<0.001, **** p<0.0001. Factorial analysis was performed using Stat-Ease 360® version 25.0.6 to evaluate the relative contribution of devitalization method, preservation strategy, and storage time on functional outcomes.

## Acknowledgments

This work was developed within the scope of the project CICECO Aveiro Institute of Materials, UID/50011/2025 (DOI 10.54499/UID/50011/2025) & LA/P/0006/2020 (DOI 10.54499/LA/P/0006/2020), financed by national funds through the FCT/MCTES (PIDDAC). It was supported by Programa Operacional Competitividade e Internacionalização, in the component FEDER, and by national funds (OE), through FCT/MCTES in the scope of the project ‘CellFi – Advancing the Regenerative and Translational Potential of Cellular Fibers’, PTDC/BTM-ORG/3215/2020. LEO Foundation funded this research in the scope of the project ‘Lazarus – Exploring devitalized MSCs fibres as immunomodulatory devices for wound regeneration’, LF-OC-23-001396. The authors also acknowledge the RoosterBio Development Award 2024. M.C.C. acknowledges the PhD grant 2024.04113.BD (DOI 10.54499/2024.04113.BD). M.B.O. acknowledges national funds through FCT – Fundação para a Ciência e a Tecnologia, I.P., under the Scientific Employment Stimulus – Institutional Call – CEECINST/00013/2021. The authors thank João Gomes for contributions to preliminary studies within his Bachelor’s project.

