## Supporting Information for "Time-Evolving Immunomodulatory Activity in Green-Fixed Devitalized Cellular Materials"

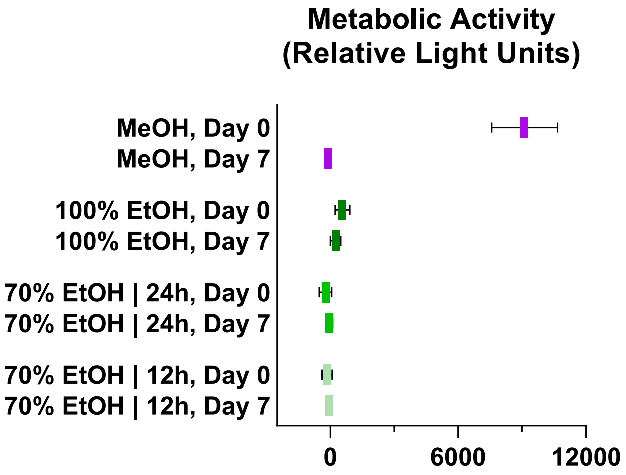


**Figure S1.** CellTiter-Glo® of MeOH and EtOH conditions after 7 days in full media.

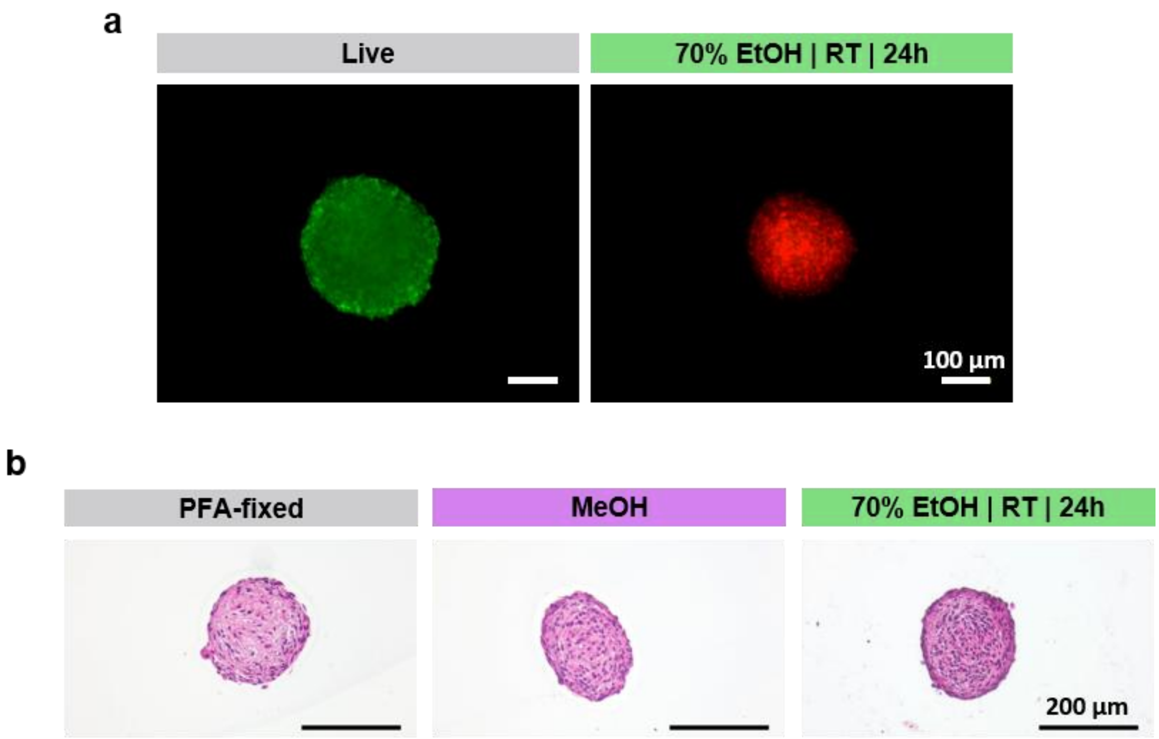


**Figure S2. a)** Live/dead staining of live vs EtOH-devitalized spheroid. Scale bars: 100 µm. **b)** H&E staining of PFA-fixed, MeOH- and EtOH-treated spheroid sections. Scale bars: 200 µm.

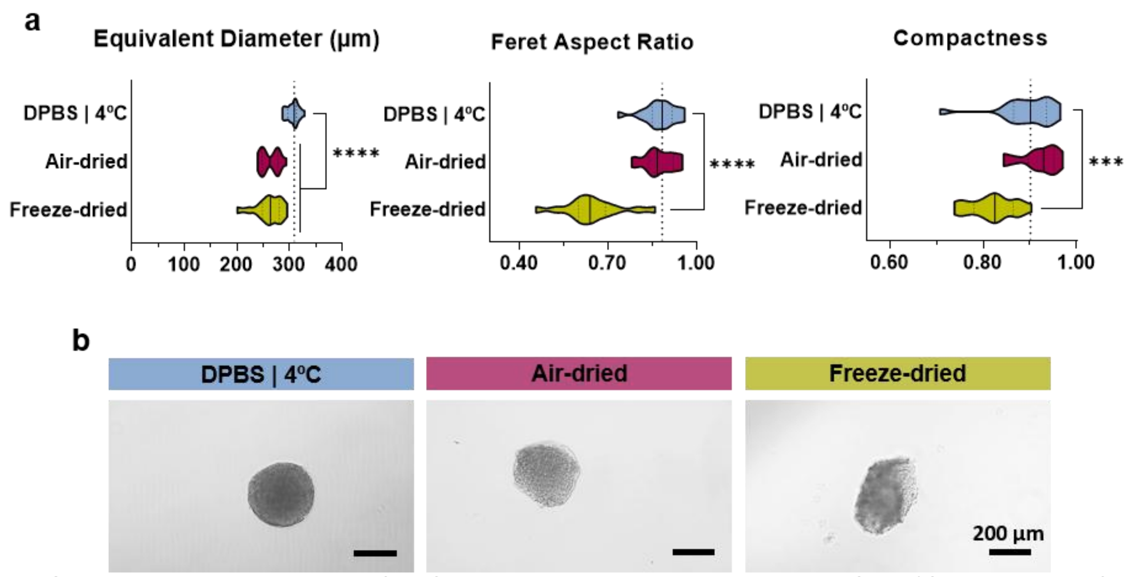


**Figure S3. a)** Morphometric analysis of MeOH-devitalized spheroids stored in DPBS at 4ºC, air-dried and freeze-dried. **b)** Representative brightfield images. Scale bars: 200 µm.

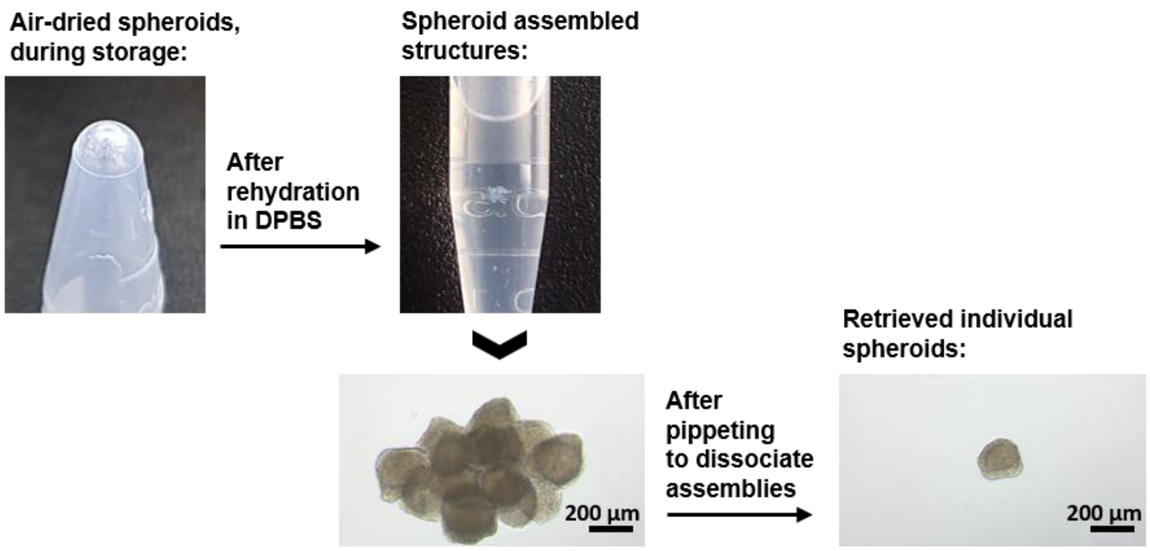


**Figure S4.** Air-dried spheroids: rehydration and handling to separate spheroid assemblies. Scale bars: 200 µm.

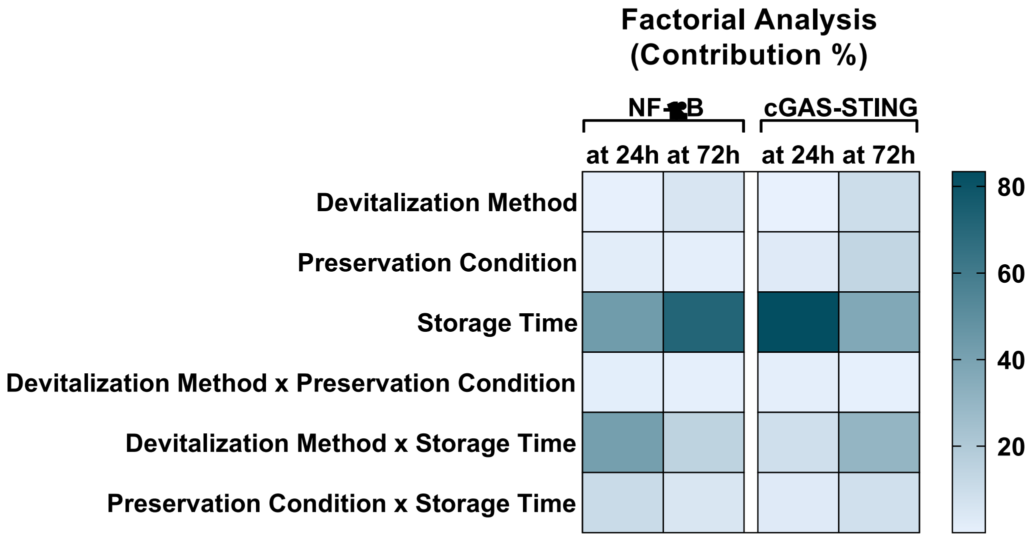


**Figure S5.** Factorial analysis shows that storage time shows a dominant contribution to the induction of changes in the immunomodulatory ability of the fixed constructs.
